# No Effect of Value on the Task Irrelevant Auditory Mismatch Negativity

**DOI:** 10.1101/2024.11.12.623328

**Authors:** Brendan T. Hutchinson, Bradley N. Jack, Alycia Budd, Ryan Calabro, Danielle Fogarty, Michael E.R. Nicholls, Oren Griffiths

## Abstract

Behavioural and neuroscientific evidence suggests visual stimuli that signal value involuntarily capture attention and are preferentially processed, even when unattended. We examined whether learned value associations for task-irrelevant auditory stimuli modulate pre-attentive processing and involuntarily capture attention. Across two experiments, the effect of learned value on the visual- and auditory-evoked mismatch negativity (MMN) and P3a event-related potential (ERP) components was measured. Participants performed a primary visual detection task while an irrelevant, unattended oddball stimulus stream was concurrently presented. Deviants within this oddball stream had been previously learned to signal one of several value outcomes: monetary reward, loss or no change. Neither the auditory nor the visual MMN was influenced by these value associations. However, stimulus value affected performance on the primary task and the magnitude of the P3a in those who could identify the stimulus-value pairings at test. Supplementary mass univariate analyses and time frequency decomposition (theta phase-locking) confirmed the presence of the MMN and the absence of any influence of stimulus value on the MMN response. Findings suggest that learned value associations do not meaningfully influence the MMN prediction signaling mechanism for task-irrelevant auditory stimuli.

## 1. Introduction

Stimuli known to signal value are preferentially processed and involuntarily capture attention. Involuntary or counterproductive attentional capture occurs when the strategic control of attention is placed in direct opposition to the influence of reward (Anderson et al., 2011; Anderson & Yantis, 2013). For example, when task-irrelevant distractors are conditioned to signal reward, their presentation slows reaction times to concurrent targets (Andersen, 2016; Anderson et al., 2011; Munneke et al., 2015). Such interference effects afford the most direct quantification of the bottom-up influence of reward on attention, as the effect of value is counterproductive and thus cannot be readily attributed to intentional, strategic responding.

Precisely how early in the processing stream involuntary selection or “attentional capture” occurs remains an open question, but evidently it occurs rapidly. Several studies show oculomotor capture by distractor stimuli that signal reward availability in the visual domain (Pearson et al., 2022). In a common variant of these tasks (Le Pelley et al., 2015), participants were required to detect an oddball amongst five distractors. On some trials, one of the distractors was coloured and signalled value. On these trials, a high reward was obtainable for a correct response to the oddball. Because the task was designed to reward speeded responding, orienting to the value-paired coloured stimulus was counterproductive, as it would either slow responding or cancel the reward delivery altogether. Nonetheless, participants’ overt attention was captured by the reward-paired stimulus. Subsequent work demonstrated that explicit attentional capture also occurs for non-salient stimuli (Failing et al., 2015), and that capture occurs rapidly, often during the first saccade (Pearson et al., 2016).

The effect of reward history on involuntary attentional capture in the auditory domain is less well characterized (Dalton & Hughes, 2014). There is behavioural evidence that associating reward with auditory stimuli can lead to involuntary attentional capture. Auditory stimuli that have been learned to signal reward worsen task performance on concurrent auditory (Asutay & Västfäll, 2016; Kim et al., 2021) and visual tasks (Anderson, 2016). Electrophysiological data suggests that the involuntary capture of attention can occur for auditory stimuli that signal reward, as these stimuli elicit larger early, primarily perceptual, event-related potential (ERP) components. Across three experiments, Folyi and colleagues (Folyi et al., 2016; Folyi & Wentura, 2019) presented participants with auditory tones that were learned to signal either monetary gain or monetary loss. Participants performed a demanding auditory discrimination task whilst these value-signalling tones were simultaneously presented. Despite being irrelevant to participants’ primary task, tones that signalled monetary gains or losses elicited larger N1 responses than neutral tones.

Recent work by Demeter et al. (2022) examined whether the effect of stimulus value extended to the mismatch negativity (MMN), an early ERP component elicited in response to a discernible change in sensory stimulation that is thought to reflect automatic pre-attentive processes (Näätänen et al., 2007). The MMN is interesting in this context because it is not merely sensory and instead requires a distinction to be drawn between the evoking stimulus and those that preceded it. Participants performed an oddball task with auditory tones that signalled either high or low reward and were to respond to these oddball tones. Compared with low-reward targets, high-reward targets elicited larger N1, MMN, and P3b (Demeter et al., 2022). Since the N1 and MMN index early stages of processing, modulation of the MMN by learned value suggests that reward associations can affect prioritisation at very early processing stages. However, stimuli that evoked an MMN response in their study were targets in an ongoing behavioural task. We can therefore expect that they were already subject to sustained endogenous attention. Under some taxonomies, their observed MMN might therefore better be understood as an N2b response (Fitzgerald & Todd, 2020).

More generally, when the effect of attention on early ERP components is studied, the modulation of attention typically occurs in paradigms where attention refers to effortful, endogenous attention to task-relevant stimuli (Luck & Kappenman, 2012). It therefore remains unclear whether these same value-modulations of the MMN response would occur for stimuli that were task irrelevant. Interestingly, while the P3a follows momentarily after the MMN, it can be functionally distinguished from the MMN. The P3a indexes the capture of explicit attention (Sussman et al., 2003). Like the MMN, the P3a is commonly studied for task irrelevant stimuli and is distinguished from the P3b on these grounds (Polich, 2007). While it is already known that the availability of reward modifies the P3b to task relevant stimuli (Hömberg et al., 1981; Yeung & Sanfey, 2004; Demeter et al., 2022), it may be possible that value-signalling task *irrelevant* stimuli modify the size of the P3a.

Across two experiments, we trained participants to learn associations between stimuli (tones or checkerboards) and monetary value (either gains or losses). These stimuli then appeared infrequently as “deviants” in an unattended stream of repeated (standard) stimuli, which was presented while participants simultaneously performed a demanding vigilance task (modelled on Winkler et al, 2005). Deviants paired with value were compared against control deviants that were not paired with value. Experiment 1 examined the effect of pairing deviant stimuli with monetary reward (relative to a non-rewarded control) on the auditory and visual MMN and P3a components. Experiment 2 exclusively focused on the auditory MMN/P3a, increased reward magnitude, and explored the impact of pairing stimuli with monetary rewards and losses. Given that learned value associations can influence both early sensory (Folyi & Wentura, 2019) and later “cognitive” processing stages (Demeter et al., 2022) for task relevant stimuli, we predicted that value-signalling stimuli would influence the magnitude of the MMN and P3a to task-irrelevant stimuli.

Supplementary to our conventional ERP analysis, both experiments also explored intertrial coherence (ITC or “phase locking”) in the theta band (4-7Hz). Phase-resetting (and the consequent increase in phase alignment) of theta frequency occurs when a stimulus is compared against other recently presented stimuli (Klimesch, 1999; Rizzuto et al., 2003), including to an auditory deviant in an oddball stimulus stream (Ko et al., 2012; Fuentemilla et al., 2008). Initial evidence came from Fuentemilla and colleagues (2008), who found that the auditory MMN within an oddball task was associated with increased theta power and phase synchronization. Broadly similar changes in theta phase locking have also been seen in visual MMN paradigms (Stothart & Kazanina, 2013; Yan et al., 2017), suggestive that theta oscillatory activity is crucial in the processing of deviants (Klimesch, 1999; Ko et al., 2009; Rizzuto et al., 2003). The increase in theta phase locking following a deviant is reliably disrupted in people with schizophrenia in auditory oddball tasks (Hochberger et al., 2019) and in other auditory tasks (Griffiths et al., 2022) and has been proposed as a biomarker of generalized theta dysfunction in schizophrenia (Javitt et al., 2018). Contemporaneously, evoked theta activity has been associated with transforming the experience of unanticipated value (i.e., a valenced prediction error) into behavioural change (Cavanagh et al., 2010), a capacity that is also disrupted in schizophrenia (Juckel et al., 2006). Thus, we additionally sought to explore whether theta responses following a deviant stimulus were modulated by associative value.

In anticipation of our primary findings, we saw little effect of value on the MMN. To ensure our a priori focus on the standard spatiotemporal profile of the MMN did not obscure our capacity to detect any unexpected effects, we additionally performed an exploratory mass-univariate analysis. This analysis corroborated our initial null finding.

## 2. Experiment 1

In Experiment 1, a task-irrelevant oddball stimulus stream with infrequent deviants that had been associated with monetary reward or no change in monetary value was presented while participants performed a primary visual task (Winkler et al., 2005). We included both an auditory and a visual variant of the oddball stimulus stream, thus enabling separate assessment of the effect of value on the MMN in both modalities. In the auditory oddball task, standards and deviants were sinusoidal auditory tones that varied only in pitch (frequency). In the visual oddball task, standards and deviants were checkerboard patterns closely matched in all respects except spatial frequency. The modality of the oddball stream varied between experimental blocks, while deviant value varied within each experimental block. The concurrent vigilance task was always visual but, in the case of the visual oddball task, the vigilance task was presented in a different vertical hemifield to the oddball stream. We hypothesized that reward-history of task irrelevant stimuli would modulate early pre-attentive change-detection processes, as well as the degree of involuntary attentional capture. These effects were predicted to be observed in a larger MMN and larger P3a, respectively, for deviants that signalled reward compared with neutral deviants.

### 2.1. Method

#### 2.1.1. Participants

Sample size was based on power analysis conducted in alignment with effect sizes derived from Demeter et al. (2022) and Folyi and Wentura (2019). The former observed an effect size for N1/MMN modulations by reward to task relevant auditory stimuli of *d* = .48, while the latter reported an effect size for N1 to task irrelevant auditory stimuli based on valence of η_p_^2^ = .145. With alpha set at .05 and power set to 0.8, these effect size estimates set sample size requirements at a conservative lower bound of 26 and an upper bound of 37. Our final sample size of 45 undergraduate psychology students for Experiment 1 (*M* age = 20.84 years, *SD* = 5.82; 20 males and 25 females; all enrolled at Flinders University) therefore ensured we were well powered to detect an effect. Participants in both Experiment 1 and Experiment 2 received course credit and $12 AUD over the course of the experiment. Both experiments were approved by Flinders University’s Social and Behavioural Ethics Committee and was conducted in accordance with the ethical standards laid down in the Declaration of Helsinki (World Medical Association, 2004).

#### 2.1.2. Materials and Apparatus

Both Experiment 1 and Experiment 2 were programmed and presented using Psychtoolbox for MATLAB (Kleiner et al., 2007) and were displayed on a 24” computer monitor running at 1920 x 1080 resolution at 60 Hz refresh rate (model number: AW2521HF). Viewing distance was approx. 60 cm. Auditory stimuli were delivered using over-ear headphones (Sennheiser HD201). All standard and deviant tones were three pure sine waves, one standard and two equidistant deviants. For Experiment 1, the standard was 1000 Hz with deviants at 900 Hz and 1100 Hz (Näätänen et al., 1978; Näätänen et al., 1982; Woldorff & Hillyard, 1991). For Experiment 2, standard and deviant tones were divided into two sets of three stimuli: one standard at 600 Hz (with deviants at 450 Hz and 750 Hz), and one standard at 1400 Hz (with deviants at 1250 Hz and 1550 Hz). Tones were measured at 80dB sound pressure level, using a handheld acoustic meter. For the visual oddball task (Experiment 1 only), black/white checkerboard patterns were presented on a black background, separated into three stimuli (one standard and two deviants). Each checkerboard had the same overall size (400 x 400 pixels, 10.10° visual angle) but differed in spatial frequency. Relative to the standard (20 x 20 checkerboard pattern), one deviant’s spatial frequency was increased (40 x 40 checkerboard), and the other deviant’s spatial frequency was decreased (10 x 10 checkerboard). A grey (RGB: 127, 127, 127) fixation cross (100 pixels, 2.53° visual angle) was presented in the horizontal centre of the upper half of the screen in between the presentation of the checkerboard patterns.

Across both experiments, deviants were associated with some value—either the delivery of reward (+ $0.02) or no change in monetary tally (Experiment 1); or delivery of reward (+ $0.10), punishment (-$0.10), or no change in monetary tally (Experiment 2). Monetary icons used for the Pavlovian procedure in which participants learned the stimulus-value pairings were 400 pixels square (9.19 x 9.68 visual degrees) that were presented in the centre of the visual display (see Figure 1).

**Figure 1.**
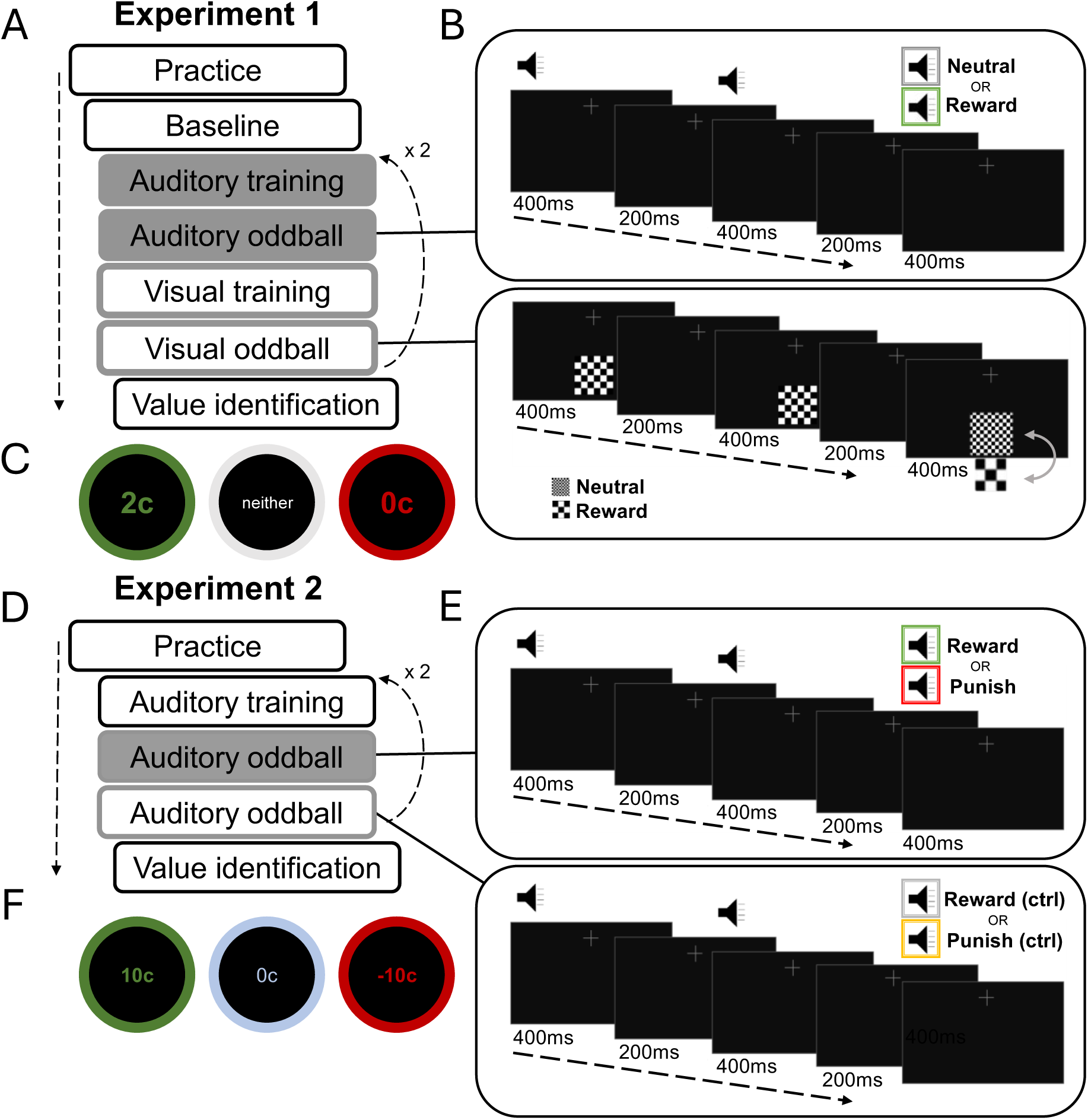
Overview of auditory / visual (Experiment 1, **A-C**) and valued (grey, unshaded) / non-valued (grey, shaded) (Experiment 2, **D-F**) oddball blocks (counterbalanced between participants). **Panel B** shows auditory task (upper) and visual task (lower). For auditory oddball task, tones (standards and deviants) were presented simultaneous with vigilance task (100ms duration, 500ms ISI). For visual oddball task, checkerboards (standards and deviants) were presented simultaneous and with identical duration as vigilance task cross (400ms duration, 200ms ISI). One deviant was presented after every 4-6 standards (here shown after 2 for illustration purposes). **C, F)** Icons (not to scale) used in training and value identification phase of experiment. **E)** Example of valued oddball sequence (upper box, grey and unshaded) and non-valued oddball sequence (lower box, grey and shaded). Tones (standards and deviants) were presented simultaneous with the presentation of the vigilance task cross (100ms duration, 500ms ISI). One deviant was presented after every 4-6 standards. In each experiment, participants completed both sets of blocks twice.

#### 2.1.3. Procedure

An overview of the general procedure is illustrated in Figure 1. Participants first performed a brief (< 30s) practice phase in which they familiarised themselves with the attentionally-demanding visual detection task (hereafter “vigilance” task). This task was adapted from Winkler and colleagues (2005) and required participants to monitor a grey cross in the upper half of the visual display, while the task irrelevant oddball stimulus stream was concurrently presented. The constantly presented cross randomly shortened by 5% on either the vertical or horizontal axis on 20% of occasions in which a stimulus appeared in the concurrent oddball stream (10% change in width: 10% change in height). This randomization had one constraint: the change never occurred during a deviant stimulus or during the standard that immediately preceded it. The cross remained shortened until a keypress was registered or until 600ms had elapsed. Responses were recorded with the space key. Participants were instructed that the vigilance task was their primary task, and to ignore any other stimuli shown or played.

##### Baseline block

Following practice, participants performed the vigilance task while being sequentially exposed to all oddball stimuli used in the experiment (prior to these stimuli being associated with value). Participants were played 120 repetitions of each of the three tones used in the auditory oddball task for a duration of 100ms and a 500ms inter-stimulus interval (ISI). Auditory stimulus duration and ISI were fixed across all phases in Experiments 1 and 2. Participants were then shown the three checkerboard stimuli used in the visual oddball task. Each checkerboard was presented 120 times (400ms duration, 200ms ISI; fixed across all phases). The order of presentation (tones first or checkerboards first) was counterbalanced between participants, but within each modality the standard stimulus was always shown first. The cross did not change in the first 3s of the Baseline block.

##### Auditory/Visual Training block

Participants passively viewed stimuli to train the stimulus-value pairings that were later presented during the main experimental blocks (see Figure 1). The two deviant stimuli (tones in auditory training, checkerboards in visual training) were shown in a random order. Assignment of deviants to value conditions was counterbalanced. One tone/checkerboard always preceded reward (e.g., 900 Hz / lower spatial frequency) while the other (e.g., 1100 Hz / higher spatial frequency) always preceded the neutral icon, and no change in tally (the other half of participants received the opposite assignment). Monetary icons were presented centrally and remained onscreen for 500ms, with a 1s inter-trial interval (ITI) between the presentation of a monetary icon and the subsequent stimulus. This continued for 40 trials (20 repetitions per conditioned stimulus, consistent with other studies of evaluative conditioning, Lipp et al., 2020; Gast & Rothermund, 2011). Participants were shown their running tally at the end of each block.

##### Auditory/Visual Oddball block

Following the training block, participants completed two oddball blocks. Visual training preceded the visual oddball blocks, and auditory training immediately preceded the auditory oddball blocks. In each oddball block, a standard was continuously repeated, and one of the two deviants occurred on every fifth, sixth or seventh presentation (amounting to a probability ratio of approximately 85% standards to 15% deviants). Each block (auditory/visual) contained 60 presentations of each deviant (neutral, reward), resulting in the presentation of 720 stimuli in total. Participants completed the vigilance task described previously while standards/deviants (tones in auditory blocks, checkerboards in visual blocks) were presented. The cross never changed size during the presentation of a deviant, or the standard that immediately preceded a deviant. Participants were not shown monetary icons. They were told that money accumulated in the same manner as during training. Participants were shown their mean response speed for the vigilance task and their running monetary tally at the end of each block. After completing the training and oddball blocks in both modalities, blocks were repeated one more time in the same order.

##### Value Identification Task

Participants finished by completing a manipulation check to assess whether they had learned the values assigned to each deviant. Each deviant was presented twice in a random order. After each stimulus was presented, two value icons (reward, neutral) were shown onscreen alongside a third response option to indicate that the participant was unsure (see Figure 1). Responses were untimed and made using the mouse. There was a 1s ITI after each response. Accuracy in this task was calculated for every stimulus type (rewarded, neutral, standard) as the proportion of reward-responses chosen for that stimulus. Participants were categorised based on their performance on the value identification task. Participants who were unable to correctly identify the reward response on both occasions were classified as “unaware”, while those who were able to correctly identify the reward response on both occasions were classified as “aware”. Twenty-nine participants were classified as aware for the auditory stimuli (*N* = 15 for unaware) and 33 participants were classified as aware for visual stimuli (*N* = 11 for unaware).

#### 2.1.4. EEG Recording and Analysis

A 32-channel BioSemi Active Two system (www.biosemi.nl) was used to record EEG data at a sample rate of 2048 Hz. Additional external electrodes were applied on the nose, below the left eye, the left and right temples, and the left and right mastoids. EEG-activity was time locked to each of the auditory tones and checkerboard stimuli that were presented throughout the duration of the experiment.

All pre-processing was completed using custom Matlab scripts written using EEGlab (Delorme & Makeig, 2004). Raw data were down sampled to 256 Hz. Then the PREP-pipeline (Bigdely-Shamlo et al., 2015) was used with default values to detrend the data, identify and interpolate noisy channels, and apply a robust median reference. Data were then band-pass filtered from 0.1 to 30 Hz using a fourth order, zero-phase Butterworth filter, and re-referenced to the mastoids. Blinks and eye movements were automatically identified and removed via independent components analysis (ICA) using the EEGlab toolbox plugin IClabel (Pion-Tonachini et al., 2019), which uses a weighted convolutional neural network to estimate the classification of components. Components classified as eye-generated with greater than 50% probability were removed. Data were epoched into 600ms time windows (200ms pre-stimulus, 400ms post-stimulus) surrounding each time-locked event and, after averaging, ERPs were baseline corrected using mean amplitude of 100ms preceding stimulus onset.

Electrophysiological data analyses were conducted using custom Matlab scripts written for EEGlab (Delorme & Makeig, 2004). For ERP analysis, electrode channels for each component (MMN, P3a) were selected based on their typical spatial profile (Näätänen, 1995). A group of frontal-central channels (Fz, F3, F4, FC1, FC2, Cz, C3, C4) were used for the MMN and a selection of central-parietal channels (CP1, CP2, P3, Pz, P4, PO3, PO4) were used for the P3a (see Polich, 2007). Components were measured based on the response to standards/deviants within the oddball experiment blocks. The primary dependent variable was mean amplitude of difference waves, calculated as the deviant minus standard (excluding standards that immediately followed deviants to avoid anticipation/expectation effects on the ERP). Mean amplitude over a 100ms time window surrounding the peak of each component was computed. As recommended by Luck and Gaspelin (2017), to ensure our time window for analysis was orthogonal to the effect(s) of interest, we first averaged together difference waves across all groups and conditions for each component separately, and then identified the most negative voltage point as the peak of the MMN and the most positive voltage point as the peak of the P3a. This was performed for the auditory and visual tasks separately, yielding the following windows: 113–213ms for the auditory MMN, 102–202ms for the visual MMN, 207–307ms for the auditory P3a, and 168–268ms for the visual P3a.

In addition to our conventional statistical analysis, a mass univariate analysis was performed with cluster mass permutation (Maris & Oostenveld, 2007) using the mass univariate analysis ERP toolbox available for Matlab (Groppe et al., 2011). Amplitude values for difference waves (down sampled to 128 Hz) from all scalp channels at all post-stimulus time points were submitted comparing rewarded deviants with neutral deviants, separately for the auditory and visual oddball task.

Quantification of theta coherence was performed on single trial data on the same epochs as the ERP analyses using EEGlab’s *newtimef* function. Default values were applied with additional padding (pad ratio = 8) and a larger window size (348ms, 89 samples) to obtain measurements of lower frequencies in the available epoch. Time-frequency amplitude estimates for 118 linearly spaced frequencies from 3.2 Hz to 50 Hz were extracted using Morlet wavelets, with the number of wavelet cycles linearly scaled from 1 to 7.8 cycles. Phase coherence was quantified as the amplitude normalized mean phase-angle over the same electrode pool and temporal window used to quantify MMN magnitude.

#### 2.1.5. Statistical Analysis

Statistical analyses were conducted using R version 4.2.0 (R Core Team, 2021) and JASP version 0.14.1 (JASP team, 2022). Both frequentist and Bayesian statistics are reported for all analyses (Keysers et al., 2020). All two-sample *t* tests used the Welch modification to the degrees of freedom. Sphericity violations within ANOVA’s were dealt with using the Greenhouse-Geisser correction to the degrees of freedom. Effect sizes are reported as Cohen’s *d* or partial-eta squared (η_p_^2^) and used small sample corrections where appropriate. Bayes factors (BF) were computed using Rouder’s method with the default Cauchy prior of 0.707 for t-tests, r = 0.5 for fixed effects, and r = 1 for random effects in ANOVAs (Morey & Rouder, 2011; Rouder et al., 2012).

### 2.2. Results

#### 2.2.1. Behavioural Performance

We first assessed whether behavioural performance on the vigilance task varied between the auditory and visual oddball task, and/or the value association of the deviants (those that had most recently occurred prior to any given task-relevant change in the vigilance task). We focused on accuracy as reaction times could only be calculated prior to the relatively tight response deadline (600ms), which led to bias within the observed scores. Accuracy data on the vigilance task was examined with a 2 (modality: auditory, visual) by 2 (deviant value: rewarded, neutral) repeated measures ANOVA. Results indicated that there was a main effect of modality on accuracy, *F*(1, 44) = 118.02, *p* < .0001, BF_10_ > 100, η_p_^2^ = 0.73, but no main effect of deviant value, *F*(1,44) = 0.50, *p* = .48, BF_10_ = 0.22, η_p_^2^ = 0.01, nor was there an interaction between modality and deviant value, *F*(1,44) = 0.49, *p* = .49, BF_10_ = 0.27, η_p_^2^ = .01. Participants made more errors (*M*_error_ = 52.13, *SD* = 20.31) during the visual oddball task than the auditory oddball task (*M*_error_ = 24.07, *SD* = 12.41), but deviant value did not appear to impact behavioural performance (see Figure 2).

**Figure 2.**
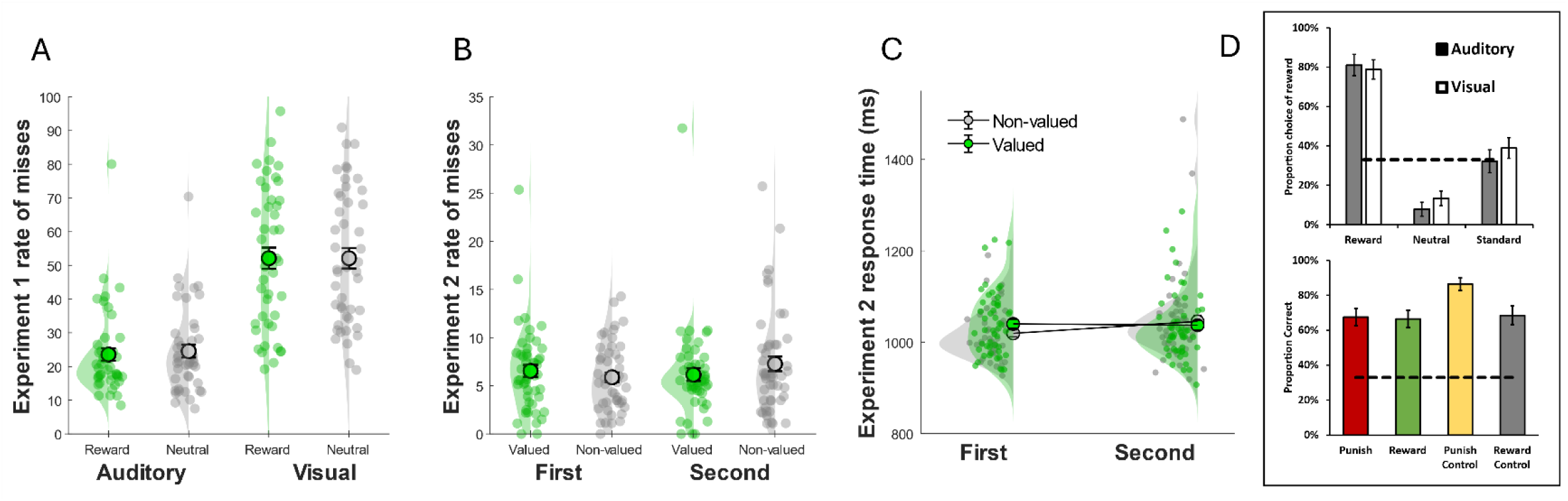
Task performance on the primary “vigilance” task. **A)** Mean error (proportion of misses) in Experiment 1 (green: rewarded deviants, grey: neutral deviants). Stimulus (reward, neutral) refers to the most recently presented deviant. **B)** Mean error (proportion of misses) in Experiment 2 (green: valued deviant block, grey: non-valued deviant block). Order (First, Second) refers to counterbalancing condition (whether participant was first exposed to valued deviant block or non-valued deviant block). **C)** Response times (ms) on the primary vigilance task by Value (Valued, Non-valued) and Order (First, Second). Insets of panel **D** show performance on the post-experiment value identification task (top: Experiment 1, bottom: Experiment 2). Experiment 1 performance indicates proportion of reward response chosen for each stimulus type. Experiment 2 performance indicates proportion correct for each stimulus type. Raw numbers for each are presented in Supplementary Table 1 and 2. Dashed lines indicate chance performance.

Accuracy on the follow-up value identification task was assessed using a 2 (modality: auditory, visual) by 3 (stimulus type: rewarded, neutral, standard) repeated measures ANOVA. Results indicated that there was no main effect of modality on test performance, *F*(1, 44) = 0.80, *p* = .38, BF_10_ = 0.22, η_p_ = .02, nor was there an interaction between modality and stimulus type, *F*(2,88) = 0.49, *p* = .62, BF_10_ = 0.13, η_p_^2^ = .01. Test performance was impacted by stimulus type, *F*(1.71, 75.39) = 89.91, *p* < .0001, BF_10_ > 100, η_p_^2^ = .67. Because our accuracy outcome measure was the mean proportion of reward response chosen, we expected higher values for rewarded stimuli and lower values for standard/neutral stimuli. This is precisely what was observed: participants responded with a high proportion of reward responses for rewarded stimuli (*M* = 80%), while proportion of reward responses for standard stimuli was approximately at chance (*M* = 35.56%) and were well below chance for unrewarded stimuli (*M* = 10.56%) (see Figure 2B).

#### 2.2.2. Electrophysiological Results

On average, 4.7 channels were interpolated, 4.9 % of trials were excluded, and 2.3 independent components were removed per participant. Average trial count per participant was 918 for standards and 114 for deviants. Difference waves can be seen in Figure 3 (raw ERPs are presented in supplementary figure 1).

**Figure 3.**
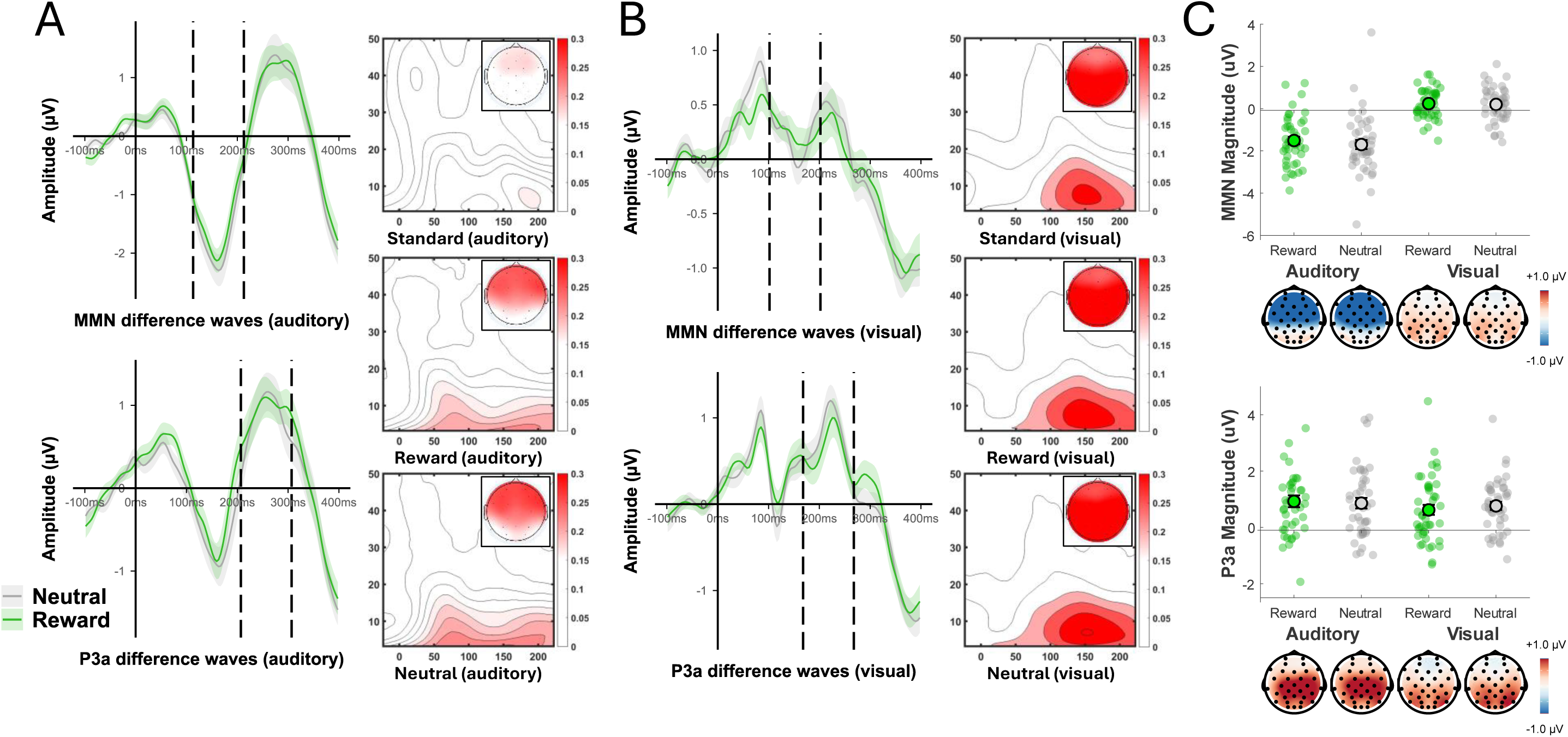
Electrophysiological results for auditory and visual oddball tasks in Experiment 1. **Panel A** shows MMN difference waves (top), P3a difference waves (bottom), and intertrial coherence (time frequency plots) for auditory oddball block. Panel B shows MMN difference waves (top), P3a difference waves (bottom), and intertrial coherence (time frequency plots) for visual oddball block. MMN difference waves were calculated over frontal-central electrodes (Fz, F3, F4, FC1, FC2, Cz, C3, C4) used to quantify the MMN. P3a difference waves were calculated over the central-parietal electrodes (CP1, CP2, Pz, P3, P4, PO3, PO4) used to quantify the P3a. Dashed vertical lines over difference waves indicate the critical time window used for statistical analysis. Shaded region around difference waves reflects +/- 1 standard error. Time frequency plots show intertrial coherence (phase alignment across trials) ranging from 0 to 1, distributed across time (x-axis) and frequency (y-axis) (top: Standard, middle: Reward, bottom: Neutral). Scalp maps in these plots indicate mean coherence at each channel averaged over theta frequency (4–7 Hz) during critical time window selected for that experimental block (refer to methods). **Panel C** shows mean amplitude of MMN difference waves (top) and P3a difference waves (bottom) during critical time window for auditory (filled) and visual (unfilled) oddball tasks. Scalp maps reflect amplitude topography over the critical time window for each ERP component. Error bars reflect +/- 1 standard error.

##### Auditory MMN

Separate one sample t-tests using mean amplitude for each condition confirmed the elicitation of an auditory MMN for the rewarded deviant, *t*(43) = 8.50, *p* < .0001, *d* = 1.28, BF_10_ > 100, and neutral deviant, *t*(43) = 7.99, *p* < .0001, *d* = 1.20, BF_10_ >100. A paired-samples t-test indicated the effect of value (reward, neutral) was not significant on the auditory MMN, *t*(43) = 1.11, *p* = .27, *d* = 0.17, BF_10_ = 0.29. Figure 3 shows a prototypical MMN effect of comparable magnitude across the two value conditions.

##### Auditory P3a

One sample t-tests confirmed the elicitation of an auditory P3a for both the rewarded deviant, *t*(43) = 4.44, *p* < .0001, *d* = 0.67, BF_10_ > 100, and the neutral deviant, *t*(43) = 4.62, *p* < .0001, *d* = 0.70, BF_10_ > 100, however a paired-samples t-test indicated the difference between valued and neutral deviants was not significant, *t*(43) = 0.37, *p* = .72, *d* = 0.05, BF_10_ = 0.17.

##### Visual MMN

Separate one sample t-tests confirmed the elicitation of a visual MMN for the rewarded deviant, *t*(43) = 2.25, *p* = .03, *d* = 0.34, BF_10_ = 1.58, but not the neutral deviant, *t*(43) = 1.62, *p* = .11, *d* = 0.24, BF_10_ = 0.54. No difference in amplitude was observed between the rewarded and neutral MMN magnitudes, *t*(43) = 0.25, *p* = .80, *d* = 0.04, BF_10_ = 0.17.

##### Visual P3a

Separate one sample t-tests confirmed the elicitation of a visual P3a for both the rewarded deviant, *t*(43) = 3.51, *p* = .001, *d* = 0.53, BF_10_ = 27.79, and neutral deviant, *t*(43) = 5.34, *p* < .0001, *d* = 0.80, BF_10_ > 100, however paired-samples t-test indicated no significant difference between them, *t*(43) = 0.75, *p* = .46, *d* = 0.11, BF_10_ = 0.21.

##### Effect of contingency awareness

It is possible that any effect of value was diluted by including participants who did not adequately learn the stimulus-to-value pairings. We therefore submitted mean amplitudes of each component to separate 2 (contingency awareness: aware, unaware) by 2 (value: rewarded, neutral) mixed factorial ANOVAs with awareness in the test phase as a between-subject factor. For the MMN in the auditory task, the main effect of awareness was not significant, *F*(1,42) = 2.55, *p* = .12, BF_10_ = 0.97, η_p_^2^ = 0.06, nor was the main effect of value, *F*(1,42) = 1.68, *p* = .20, BF_10_ = 0.37, η_p_^2^ = .04, or the interaction between awareness and value, *F*(1,42) = 0.61, *p* = .44, BF_10_ = 0.56, η_p_^2^ = .01. For the P3a in the auditory task, the main effect of awareness was not significant, *F*(1,42) = 1.36, *p* = .25, BF_10_ = 0.65, η_p_^2^ = .03, nor was the main effect of value, *F*(1,42) = 0.02, *p* = .88, BF_10_ = 0.23, η_p_^2^ < .001, or the interaction between awareness and value, *F*(1,42) = 0.37, *p* = .55, BF_10_ = 0.34, η_p_^2^ = .009. For the MMN in the visual task, the main effect of awareness was not significant, *F*(1,42) = 0.41, *p* = .52, BF_10_ = 0.36, η_p_^2^ = .01, nor was the main effect of value, *F*(1,42) < .001, *p* = .98, BF_10_ = 0.23, η_p_ < .001, or the interaction between awareness and value, *F*(1,42) = 0.24, *p* = .63, BF_10_ = 0.35, η_p_^2^ = .006. For the P3a in the visual task, the main effect of awareness was not significant, *F*(1,42) = 0.38, *p* = .54, BF_10_ = 0.37, η_p_^2^ = .009, nor was the main effect of value, *F*(1,42) = 1.90, *p* = .18, BF_10_ = 0.29, η_p_^2^ = .04, or the interaction between awareness and value, *F*(1,42) = 2.09, *p* = .16, BF_10_ = 0.71, η_p_^2^ = .05.

As a further supplementary check, we drilled down more by repeating analyses within the “aware” only participants. In all cases, the effect of value was not significant: auditory MMN: *F*(1,28) = 0.17, *p* = .68, BF_10_ = .28, η_p_^2^= .006; auditory P3a: *F*(1,28) = 0.41, *p* = .53, BF_10_ = .33, η_p_^2^ = .01; visual MMN: *F*(1,32) = 0.25, *p* = .62, BF_10_ = .28, η_p_^2^ = .008; and visual P3a: *F*(1,32) = 0.005, *p* = .95, BF_10_ = 0.25, η_p_^2^ < .001. In sum, all main effects and interactions were not significant, indicating that there was no effect of value that was being ‘hidden’ based on contingency awareness.

##### Mass Univariate Analyses

To next ensure that real effects of value were not missed due to our a priori focus on the typical spatiotemporal parameters of an MMN response, mass univariate analyses were additionally applied. Separate comparisons of rewarded versus neutral difference (deviant – standard) waves resulted in no significant clusters in either the auditory (all *p* > .63) or visual task (all *p* > .47).

##### Theta coherence

Theta coherence was aggregated over frontal-central electrodes during the same temporal windows used to quantify the MMN responses. For the auditory oddball block, a repeated measures ANOVA with mean theta coherence for each stimulus (rewarded deviant, neutral deviant, standard) indicated a large main effect of stimulus, *F*(2,86) = 16.86, *p* < .0001, BF_10_ >100, η_p_^2^ = 0.28. Pairwise comparisons suggested that mean theta coherence was weaker for standards (*M* = 0.17, SD = 0.05) compared with rewarded deviants (*M* = 0.24, *SD* = 0.10) and neutral deviants (*M* = 0.25, *SD* = 0.12) (both comparisons *p* < .0001). Similarly, there was a main effect of stimulus in the visual oddball block, *F*(1.72, 73.97) = 4.90, *p* = .01, BF_10_ = 3.83, η_p_^2^ = 0.10. Pairwise comparisons suggested that mean theta coherence was significantly weaker for standards (*M* = 0.27, *SD* = 0.10) compared with neutral deviants only (*M* = 0.30, *SD* = 0.12) (*p* = .004). For completeness, difference conditions (deviant minus standard) were examined with paired-samples t-tests for the auditory and visual task separately. There was no difference in theta coherence between the reward and neutral condition in either the auditory oddball task, *t*(43) = 0.81, *p* = .42, *d* = 0.12, BF_10_ = 0.22, or the visual oddball block, *t*(43) = 0.95, *p* = .35, *d* = 0.14, BF_10_ = 0.25 (see Figure 3).

### 2.3. Discussion

Responses to auditory deviants differed in three ways from responses to standards: an early frontocentral MMN component (associated with event-related theta phase alignment), a later centroparietal P3a component, and a frontal change in theta-band phase coherence. None of these responses were affected by stimulus value. It is possible that the absence of value modulation of the MMN was due to participants simply failing to learn the stimulus-value pairings. However, this interpretation appears unlikely because (1) participants’ aggregate responses in the value identification test exceeded chance, and (2) when ERP analyses were limited to those who passed the value identification test, the same general pattern held. The results in the visual domain were more nuanced. Significant MMN and P3a effects were each nominally observed (notwithstanding the MMN following the neutral visual deviant), although their temporal windows and spatial distributions overlapped, which muddies their interpretation. Second, performance on the primary visual task was lower in the visual than auditory oddball blocks. Participants may have been orienting to the visual gratings during the visual oddball task and thus may not have treated them as task irrelevant.

We therefore shifted focus to the auditory MMN in Experiment 2. In addition to reward, here we also introduced a “punishment” condition. Stimuli that signal threat and/or punishment involuntarily capture attention in other tasks (Schmidt et al., 2015a, b) and this inclusion allowed us to explore whether reinforcement valence differentially impacts the MMN and/or P3a (Folyi & Wentura, 2019). Although other studies that have reported positive effects have used similar magnitude rewards (Anderson et al., 2011; Le Pelley et al., 2015; Theeuwes & Belopolsky, 2012), we strengthened our value manipulation by increasing the size of each reward (and punishment) by a factor of five. Finally, we organized the task/block structure such that valued stimuli were presented in one block and non-valued control stimuli were presented in a separate block. This allowed us to more directly determine whether stimulus value impacted the concurrent vigilance task by comparing behavioural performance across valued and non-valued control blocks.

## 3. Experiment 2

Experiment 2 used four distinct deviant tones and two standards (no gratings were used). Two oddball sequences were generated, each comprised of one standard tone, and two deviant tones (one with a frequency above the standard, one equidistant below). One sequence included two deviant tones that were each associated with either a gain or loss of 10c. The other sequence included two deviants that were not associated with a change in value. In this manner, a higher pitched deviant that was paired with value (reward or punishment) could be compared against a higher pitched deviant that was not (in the second, control oddball sequence). Similarly, any impact of value on the vigilance task could be observed by comparing task performance during the valued oddball sequence relative to the perceptually similar non-valued sequence. We anticipated that MMN and P3a magnitude would differ between the value-paired deviants and their matched control deviants, that punishment-paired stimuli may loom larger than reward-paired stimuli (Kahneman & Tversky, 1979), and that performance on the concurrent vigilance task would be disrupted during the value-paired oddball sequence.

### 3.1. Method

#### 3.1.1. Participants

Fifty undergraduate psychology students (*M* age = 22.94 years, *SD* = 6.82; 22 males and 28 females) from Flinders University participated.

#### 3.1.2. Procedure

The procedure was the same as in Experiment 1, with the following exceptions. Instead of switching to vision for a second training/oddball loop, participants continued with tones of a different frequency (450/600/750Hz then 1250/1400/1550Hz, or vice versa). Participants completed a single training block with all four deviant stimuli (450, 750, 1250 and 1550Hz), each presented 20 times in a random order (80 trials total). Deviants were associated with either value (-10c or +10c) or no value (0c). For example, one person might experience the two lower-pitched tones associated with value (e.g., 450Hz paired with -10c, 750Hz paired with +10c) and the remaining stimuli (1250Hz, 1550Hz) paired with no value (0c). For half of the participants, the lower frequency deviants (450, 750Hz) were paired with value; for the other half, the higher tones (1250Hz, 1550Hz) were paired with value. Crossed with this counterbalancing factor, half of the participants experienced reward following the higher pitched tone within the rewarded oddball sequence (750Hz or 1550Hz), and punishment following the relatively lower pitched deviant (450Hz or 1250Hz). This assignment was reversed for the other half of participants. Participants sequentially completed two oddball blocks with these stimuli. Presentation order of experimental blocks was counterbalanced between participants (see Figure 1). The response window for each target in the concurrent vigilance task (fixation cross detection task) was extended to 3s to allow investigation of response latencies. As previously, the target could not occur during the presentation of a standard, nor during the presentation of the standard immediately preceding a deviant.

After two repeats of training followed by two oddball blocks (valued and non-valued), participants finished with the value identification task, which differed slightly from Experiment 1: All 4 deviant stimuli were tones (450, 750, 1250 and 1550Hz) and the response options were -10c, 0c or 10c. The length of the value identification task was doubled (from two repetitions per stimulus to four). Every trial during this task was scored as correct or incorrect. Participants were considered “top learners” (n = 18) if their total score was 13 or more (∼ 80% correct) and “moderate learners” if their total score was below 13 (n = 24).

#### 3.1.3. EEG Recording and Analysis

EEG recording and analysis followed the same procedure as Experiment 1, including the same electrode groupings for each component and frontocentral theta coherence analysis, and the same “collapsed localizer” method (Luck & Gaspelin, 2017) over difference waves for identification of critical time windows. A time window of 98–198ms was used for the MMN (and theta coherence) and 219–319ms for the P3a. A mass univariate analysis was performed again to supplement our conventional statistical analysis, however here post-stimulus scalp channel time series data were submitted to a factorial mass univariate analysis (Fields, 2017) with a 2 (value: valued, non-valued) by 2 (valence: punish, reward) repeated measures ANOVA design.

### 3.2. Results

#### 3.2.1. Behavioural Results

Participants error scores (the number of changes missed within the vigilance task) and reaction times on the primary vigilance task are summarized in Figure 2. Most targets elicited a response within the 3s response window, so our primary dependent variable was response latency in Experiment 2. The error and response latency data were analysed with separate 2 (presentation order: first repetition, second repetition) by 2 (value: oddball sequence with either valued or neutral deviants) repeated measures ANOVAs. Results indicated there was a significant interaction between value and order for both error scores, *F*(1, 49) = 4.12, *p* < .05, BF_10_ = 0.12, η_p_^2^ = 0.08, and response times, *F*(1,49) = 5.82, *p* = .02, BF_10_ = 1.71, η_p_^2^ = .11. There was a significant main effect of order on response times, *F*(1, 49) = 4.05, *p* < .05, BF_10_ = 0.80, η_p_^2^ = .08, but no main effect of value on response times, *F*(1,49) = 1.35, *p* = .25, BF_10_ = 0.28, η_p_^2^ = 0.03, nor were there any main effects for error scores. Within the first oddball sequence, participants were faster to detect the visual target during the non-valued oddball sequence than the valued oddball sequence, *t*(49) = 3.09, *p* = .003, *d* = 0.44, BF_10_ = 9.86. This effect did not persist into the second repetition of each oddball sequence: valued versus non-valued, *t*(49) = 0.92, *p* = .36, *d* = 0.13, BF_10_ = 0.22. The presence of value-paired deviants distracted people from the vigilance task more than control deviants, but this diminished across training (see Figure 2D). The error data showed a broadly similar pattern of means, so there was little evidence of a speed-accuracy trade-off (see Figure 2C).

As can be seen in Figure 2E, accuracy on the value identification task was well above chance (33%), suggesting that in aggregate participants learned the stimulus-value pairings. Effects of valence and value on accuracy were assessed using a 2 (valence: reward, punish) by 2 (value: valued, non-valued) repeated measures ANOVA. This indicated that there was a main effect of valence, *F*(1,49) = 6.52, *p* = .01, BF_10_ = 1.33, η_p_^2^ = 0.12, a main effect of value, *F*(1,49) = 6.79, *p* = .01, BF_10_ = 1.93, η_p_^2^ = 0.12, and a close-to-significant significant interaction between valence and value, *F*(1,49) = 3.51, *p* = .07, BF_10_ = 5.73, η_p_^2^ = 0.07. This marginally significant interaction appeared to be driven by the fact that participants more accurately classified non-valued punished deviants than other stimuli: non-valued rewarded deviants, *t*(49) = 2.95, *p* = .005, *d* = 0.42, BF_10_ = 7.00, valued punished deviants, *t*(49) = 3.24, *p* = .002, *d* = 0.46, BF_10_ = 14.26, and valued rewarded deviants, *t*(49) = 3.88, *p* = .0003, *d* = 0.55, BF_10_ = 82.05 (see Figure 2E).

#### 3.2.2. Electrophysiological Results

On average, 6.3 channels were interpolated, 6.1 % of trials were excluded, and 2.6 independent components were removed per participant. Average trial count per participant was 929 for standards and 115 for deviants. Difference waves for each condition are shown in Figure 4 (raw ERPs are presented in supplementary figure 2).

**Figure 4.**
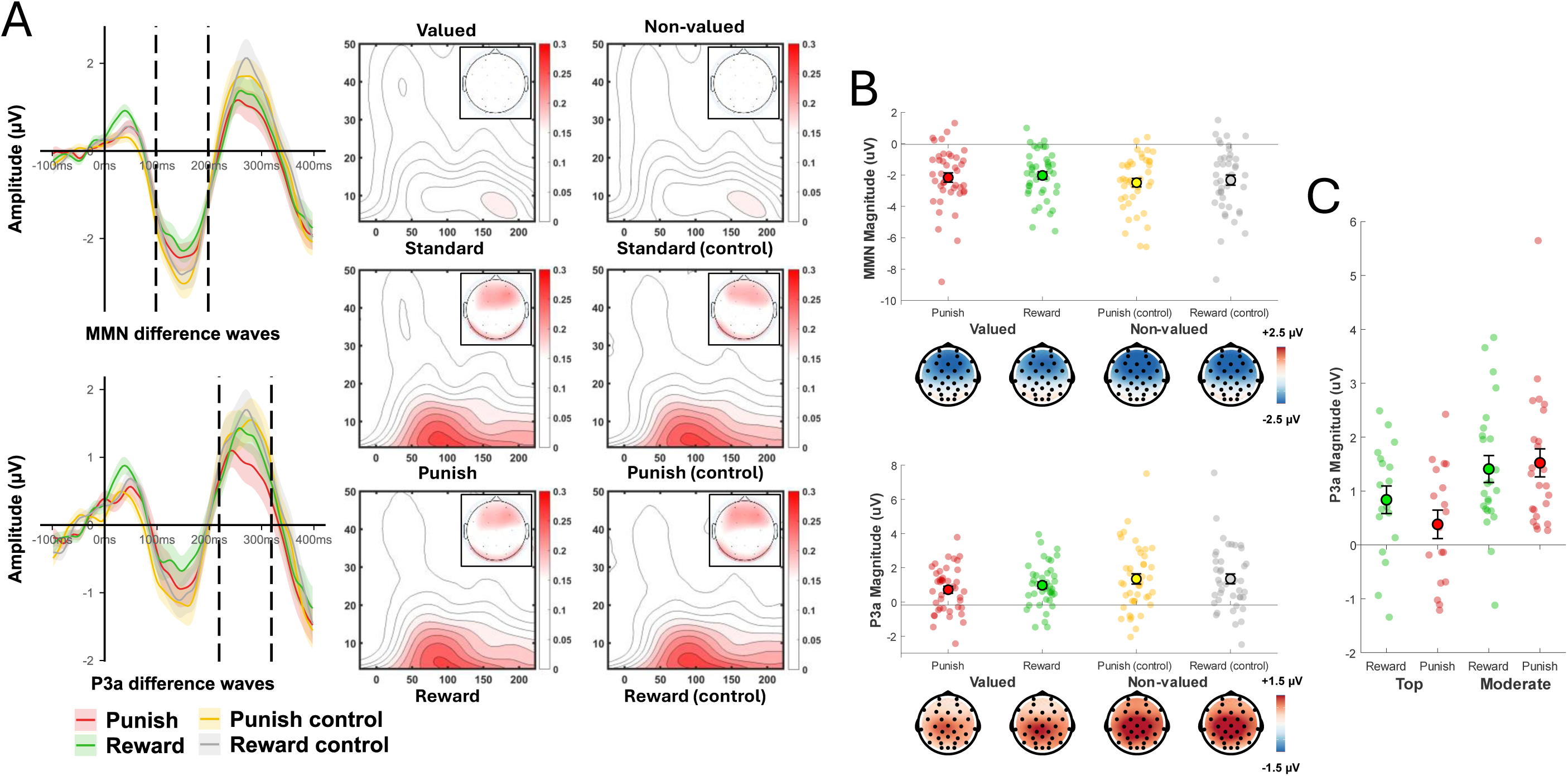
Electrophysiological results for auditory oddball tasks in Experiment 2. **Panel A** shows MMN difference waves (top), P3a difference waves (bottom), and intertrial coherence (time frequency plots) for valued and non-valued oddball blocks. MMN difference waves were calculated over frontal-central electrodes (Fz, F3, F4, FC1, FC2, Cz, C3, C4) used to quantify the MMN. P3a difference waves were calculated over the central-parietal electrodes (CP1, CP2, Pz, P3, P4, PO3, PO4) used to quantify the P3a. Dashed vertical lines over difference waves indicate the critical time window used for statistical analysis. Shaded region around difference waves reflects +/- 1 standard error. Time frequency plots show intertrial coherence (phase alignment across trials) ranging from 0 to 1, distributed across time (x-axis) and frequency (y-axis) for both the valued (left) and non-value (right) oddball blocks (top: Standard, middle: Punish, bottom: Reward). Scalp maps in these plots indicate mean coherence at each channel averaged over theta frequency (4–7 Hz) during critical time window selected for that experimental block (refer to methods). **Panel B** shows mean amplitude of MMN difference waves (top) and P3a difference waves (bottom) during critical time window for valued (filled) and non-valued (unfilled) oddball tasks. Scalp maps reflect amplitude topography over the critical time window for each ERP component. Error bars reflect +/- 1 standard error. **Panel C** shows mean amplitude of P3a difference waves grouped by contingency knowledge (top learners, moderate learners), where “top” refers to participants who classified deviants in the value identification task above 80% and “moderate” refers to participants who performed below 80%. Error bars reflect +/- 1 standard error.

##### Auditory MMN

Separate one sample t-tests performed on mean amplitude for each condition within the pre-selected time window confirmed the elicitation of the MMN: punished deviant: *t*(41) = 7.28, *p* < .0001, *d* = 1.12, BF_10_ > 100, rewarded deviant: *t*(41) = 8.52, *p* < .0001, *d* = 1.32, BF_10_ > 100, punished control deviant: *t*(41) = 9.31, *p* < .0001, *d* = 1.44, BF_10_ > 100, and rewarded control deviant: *t*(41) = 7.20, *p* < .0001, *d* = 1.11, BF_10_ > 100. A 2 (value: valued, non-valued) by 2 (valence: punish, reward) repeated measures ANOVA indicated the main effect of value was not significant, *F*(1, 41) = 1.11, *p* = .30, η_p_^2^ = .03, BF_10_ = 0.54, nor was the main effect of valence, *F*(1, 41) = 0.75, *p* = .39, η_p_^2^ = .02, BF_10_ = 0.31, nor their interaction, *F*(1,41) = 0.01, *p* = .91, η_p_^2^ < .001, BF_10_ = 0.04. Figure 4 illustrates that each deviant elicited an MMN during the critical time window, however with minimal difference in amplitude based on value or valence.

##### Auditory MMN control analysis

When value and valence stimulus assignment counterbalancing variables were added to the preceding analysis, large interaction terms were generated (see supplementary figure 3). Our counterbalancing design held linear distance between the deviants and the standards equal across conditions (all deviants were 150 Hz from their standard). However, pitch perception (stimulus frequency) is more commonly quantified non-linearly using octaves, tones and semitones. This meant that deviants in one counterbalancing condition were 2.56 semitones more distant from their standard (600 Hz standards paired with 450 Hz / 750 Hz deviants) than the other condition (1400 Hz standards paired with 1250 Hz / 1550 Hz deviants). For the second counterbalancing variable (valence), the low tone in each set (450 Hz, 1250 Hz) was 0.66 semitones more distant from the standard than the high tone (750 Hz, 1550 Hz). MMN magnitudes increase with distance from the standard, as measured in semitones (Näätänen & Winkler, 1999). Although these differences were counterbalanced across individuals, their size relative to the effect of reward led to interaction terms in the ANOVA defined by reward status.

To address this, we conducted two further analyses. First, two separate between-subjects ANOVAs were conducted to examine the effect of reward on *physically identical* stimuli. Participants were divided into three groups according to the reinforcement status of the 450 Hz and 750 Hz tones. For a quarter of the participants, the comparison between 450 Hz and 750 Hz constituted a comparison of reward versus loss. For another quarter of participants, this entailed a comparison of loss versus reward. For the other half of participants, neither deviant (450 Hz, 750 Hz) was paired with value. Figure 5A shows the mean size of this comparison for each reinforcement group separately. Most importantly, the effect of reinforcement, after controlling for physical characteristics, was not significant, *F*(2, 39) = 0.42, *p* = .66, *η_p_^2^* = .02, BF_10_ = 0.23. The equivalent analysis for the 1550 Hz / 1250 Hz comparison similarly yielded no effect of reinforcement status on the MMN magnitude: *F*(2, 39) = 1.00, *p* = .38, *η_p_^2^* = .05, BF_10_ = 0.34. Thus, no effect of reward on MMN was observed after controlling for the ratio-difference between counterbalancing conditions.

**Figure 5.**
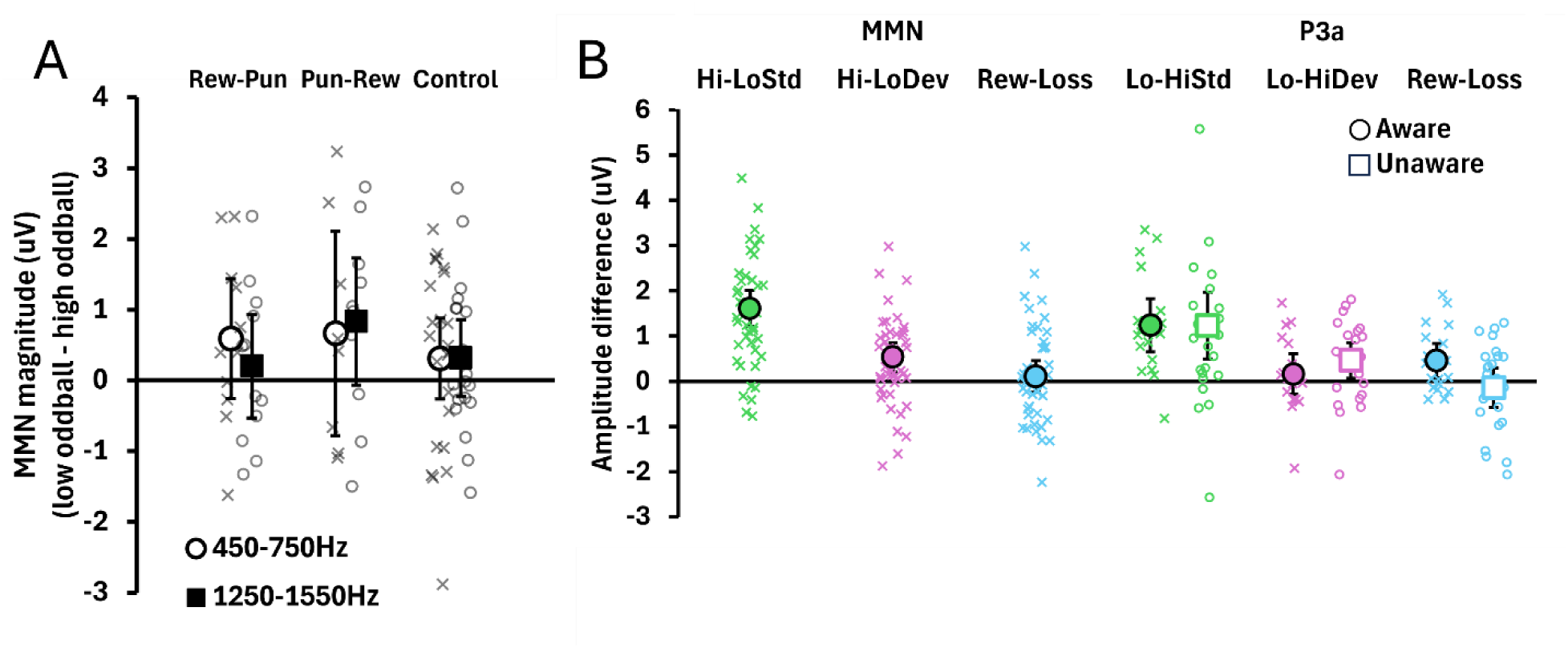
Mean differences in MMN and P3a magnitudes in Experiment 2. **Panel A** shows the effect of reinforcement on physically identical stimuli: open circles represent the 450 Hz deviant minus 750 Hz deviant, and closed circles show the difference between the 1250 Hz and 1550 Hz deviants. **Panel B** shows the mean differences for MMN and P3a magnitudes brought about by differences in the standard tone (green: 600 Hz to 1400 Hz), differences in whether the deviant represented a 150 Hz increase or decrease from the standard tone (pink) or the effect of the presence of reinforcement (blue). The P3a data, shown on the right, were further separated by whether participants showed a moderate (unaware) or high (aware) level of knowledge about the tone-value pairings. Error bars reflect +/- 1 standard error of the mean, and overlaid scatter plots represent the performance of individual participants.

Second, we quantified the size of the effect of physical deviance on MMN magnitude (controlling for reward status) to enable more direct comparisons with the magnitude of effect of reward with that of physical deviance. The first counterbalancing variable manipulated the standard tone (600, 1400Hz). The deviants paired with the 1400 Hz standard (1250 Hz, 1550 Hz) were 2.56 semitones less distant from their standard than were those paired with the 600 Hz standard (450 Hz, 750 Hz). With respect to the second counterbalancing variable, the status of a deviant as higher pitched than its matched standard (750 Hz, 1550 Hz) were 0.66 less distant from their standard than were the relatively lower pitched tones in each set (450Hz, 1250Hz). Thus, in quantifying the effect of physical deviance, we would expect the manipulation of the standard to have a larger effect on MMN magnitude than that of the deviant.

A 2 by 2 within-subjects ANOVA examined the effect of standard tone assignment (high or lower standard tone) versus the effect of deviant tone assignment (whether the deviant was relatively higher pitch or lower pitched than the standard). The main effect of standard tone assignment was significant, *F*(1, 41) = 61.80, *p* < .001, *η_p_^2^* = .60, BF_10_ > 100, as was that of the deviant tone assignment, *F*(1, 41) = 7.62, *p* = .009, *η_p_^2^* = .16, BF_10_ = 5.10, whereas the interaction was not significant, *F*(1, 41) = 0.02, *p* = 0.89, *η_p_^2^* < .001, BF_10_ = 0.22. Effect sizes (generalized eta-squared, η_G_^2^) were used to compare the magnitude of the effects of reward with those of physical stimulus differences. As shown in Figure 5B, the 2.56 semitone difference had the largest effect on MMN magnitude (*η_G_^2^* = 0.17), followed by the 0.66 semitone difference (*η_G_^2^* = 0.02). Most importantly, the presence of reward/punishment was smallest (*η_G_^2^* = 0.007) and was not statistically significant.

##### Auditory P3a

Separate one sample t tests confirmed elicitation of auditory P3a for each condition: punished deviant: *t*(41) = 3.51, *p* = .001, *d* = 0.54, BF_10_ = 27.27, rewarded deviant: *t*(41) = 5.10, *p* < .0001, *d* = 0.79, BF_10_ > 100, punished control deviant: *t*(41) = 4.54, *p* < .0001, *d* = 0.70, BF_10_ > 100, and rewarded control deviant: *t*(41) = 4.71, *p* < .0001, *d* = 0.73, BF_10_ > 100. A 2 (value: valued, non-valued) by 2 (valence: punish, reward) repeated measures ANOVA indicated there was no interaction between value and valence, *F*(1, 41) = 0.62, *p* = .44, η_p_^2^ = .02, BF_10_ = 0.09, nor was there a main effect of valence, *F*(1, 41) = 0.85, *p* = .36, η_p_^2^ = .02, BF_10_ = 0.25. However, the main effect of value was close to significant, *F*(1, 41) = 3.28, p = .077, η_p_^2^ = .07, BF_10_ = 1.09. Figure 4 illustrates that each deviant elicited a P3a, with non-significantly more positive voltage for non-valued deviants (*M* = 1.35, *SE* = 0.22) compared with valued deviants (*M* = 0.85, *SE* = 0.22).

##### Effect of contingency knowledge

To ensure that no effects were obscured based on participants’ knowledge of what each tone signalled, we submitted mean amplitudes of each component to separate 2 (contingency knowledge: top learners, moderate learners) by 2 (valence: punish, reward) by 2 (value: valued, not valued) mixed factorial ANOVAs with contingency knowledge in the test phase as a between-subject factor. The first model (MMN) indicated no significant effects (all *p* > .30). The second model (P3a) indicated that there was a significant interaction between knowledge and valence, *F*(1, 40) = 4.30, *p* = .045, η_p_^2^ = .10, BF_10_ = 0.80, a close to significant interaction between knowledge and value, *F*(1, 40) = 3.39, *p* = .07, η_p_^2^ = .08, BF_10_ = 1.38, and a main effect of knowledge, *F*(1, 40) = 6.55, *p* = .01, η_p_^2^ = .14, BF_10_ = 2.57. All other effects were not significant (all *p* > .12). The significant interaction was driven by the fact that top learners had larger P3a amplitudes following rewarded deviants than following punished deviants, *t*(35) = 2.12, *p* = .04, *d* = 0.35, BF_10_ = 3.52; whereas in moderate learners, no differences in P3a occurred between rewarded versus punished deviants, *t*(47) = 0.54, *p* = 0.59, *d* = 0.08, BF_10_ = 0.25. Value-pairing influenced P3a in those who maximally learned the stimulus-value contingencies in the value identification task (see Figure 4).

##### Mass Univariate Analyses

The mass univariate analysis used the same 2 (value: valued, non-valued) by 2 (valence: punish, reward) repeated measures ANOVA structure as our conventional ANOVA. No significant clusters were identified in the interaction or main effect of valence (all *p* > .36). However, the main effect of value showed a non-significant cluster (*p* = .11) that likely reflected the P3a effect observed in our conventional statistical analyses. The cluster spread across central-parietal and frontal scalp channels from 266 to 375ms, with a temporal peak at 312ms and spatial peak at frontal channel FP2.

##### Theta coherence

Differences in mean theta coherence (98-198ms) across the frontocentral electrode pool between each stimulus condition (rewarded deviant, punished deviant, standard) were examined with a repeated measures ANOVA separately for each block (valued, non-valued). For the valued oddball block, there was a large main effect of stimulus, *F*(1.62, 64.64) = 32.30, *p* < .0001, BF_10_ > 100, η_p_^2^ = 0.45. Pairwise comparisons suggested that mean theta coherence was weaker for standards (*M* = 0.10, SD = 0.05) compared with rewarded deviants (*M* = 0.17, *SD* = 0.07) and punished deviants (*M* = 0.18, *SD* = 0.07) (both comparisons *p* < .0001). There was also a main effect for stimulus in the non-valued oddball block, *F*(1.44, 57.74) = 25.10, *p* < .0001, BF_10_ >100, η_p_^2^ = 0.39. Pairwise comparisons suggested that mean theta coherence was weaker for standard controls (*M* = 0.10, SD = 0.05) compared with rewarded controls (*M* = 0.18, *SD* = 0.07) and punished controls (*M* = 0.17, *SD* = 0.07) (both comparisons *p* < .0001). Mean theta coherence difference values (deviant – standard) were then submitted to a 2 (value: valued, non-valued) by 2 (valence: punish, reward) repeated measures ANOVA (see Figure 4). The main effect of value was not significant, *F*(1, 40) = 0.92, *p* = .34, BF_10_ = 0.28, η_p_ ^2^= .02, nor was the main effect of valence, *F*(1, 40) = 0.20, *p* = .66, BF_10_ = 0.27, η_p_^2^ = .01, nor their interaction, *F*(1,40) = 0.92, *p* = .34, BF_10_ = 0.49, η_p_^2^= 0.02.

### 3.3. Discussion

Results generally aligned with those of Experiment 1. Robust MMN and P3a effects were observed for all deviants (valued and non-valued). Time frequency analyses again indicated stronger theta-band phase alignment following deviants than standards during the MMN measurement window. Yet, despite clear evidence that participants learned the value associations for each deviant type (refer to Figure 2E), no differences in MMN magnitude between valued and non-valued deviants was observed, either in our planned measurement window or elsewhere (assessed using mass univariate analyses). The same pattern was generally true of the P3a response, with one exception: when compared with moderate learners, top learners P3a differed between rewarded and punished deviants (but not matched control deviants). This suggests that the P3a is sensitive to stimulus value in those who most strongly learned the trained stimulus associations at test. Finally, the presence of a significant effect of stimulus value on response times during the concurrent vigilance task provided direct evidence that our stimulus value manipulation was sufficient to affect behaviour at the time that ERP responses were measured.

## 4. General discussion

The present study examined whether learned value associations involving task irrelevant stimuli modulated MMN and P3a responses. Across two experiments, participants performed a visual detection task while an oddball sequence involving value-paired deviant stimuli was simultaneously presented in an unattended secondary stream. With a total sample size of 95 participants, the MMN was not modulated by stimulus-value associations. In Experiment 1, no effect of value on MMN magnitude was observed in either the auditory or visual modality. To increase experimental sensitivity to any impact of associative value, Experiment 2 increased the value manipulation five-fold, focused exclusively on the most robust MMN effect (the auditory MMN), and altered the concurrent vigilance task to allow online measurement of behavioural distraction brought about by the oddball sequence. With these changes, we found that the associative value of stimuli in the task irrelevant oddball stream influenced performance on the primary vigilance task; and yet, once again, we saw no evidence that stimulus value affected the MMN response to task irrelevant, unattended stimuli.

### 4.1. No Effect of Value on the Auditory MMN

Our primary hypothesis concerned the effect of value on the MMN response. We reliably observed null effects, with Bayes Factors suggesting anecdotal to moderate support for each null result (BF_01_ = 1.85-5.88 across Experiments 1 and 2). The exploratory mass univariate analyses further suggested that we were not merely looking in the wrong temporal or spatial window. Of course, when faced with a null effect, it is inevitable to question the sufficiency of the manipulation and the precision of measurement of the dependent variable. Perhaps the value manipulation was too weak to induce changes in pre-attentive processes indexed by the MMN, or perhaps participants did not learn the stimulus-value mappings, or maybe our protocol was insufficient to measure modulations of the components of interest. Power calculations suggested that our experiments were sufficiently powered, but these can only serve as a rough guide when exploring a novel effect. Three additional aspects of the present data offer assurance that we were sufficiently powered to detect an effect of reward on MMN magnitude: (1) behavioural responses that demonstrated contingency awareness, (2) control analyses that showed evidence of small changes by physical deviance, and (3) associated value impacted upon behavioural responses (in Experiment 1) and P3a magnitude (in Experiment 2).

#### 4.1.1. Behavioural Evidence for Contingency Awareness

First, in both experiments, performance on the value identification task (our primary manipulation check) was strongly indicative that participants learned the trained contingencies. Because our outcome measure for this task in Experiment 1 was the proportion of reward responses chosen for each stimulus type, the response pattern observed is precisely what would be expected had participants learned the value contingencies (see Figure 2B). Similarly, in Experiment 2, where we instead examined participants’ classification accuracy for each class of deviant, response accuracies averaged 72.3% (see Figure 2E). Thus, we can be confident that participants learned the value associations. Notably, this post-test measure would - if anything - underestimate contingency knowledge, because it required participants not only to have learned the stimulus-value mappings, but to then recall those contingencies much later during test. Relatedly, in Experiment 2, when we limited our analyses to only those “top learners” who performed with greater than 80% accuracy in the value identification task (determined based on the cumulative probability of a binomial distribution, where the likelihood of a score of 13 out of 16 based on chance is *p* = .01), we still saw no effect of stimulus value on the MMN.

#### 4.1.2. Small Physical Differences Impact MMN Magnitude

Supplemental analyses within Experiment 2 reliably detected differences in MMN magnitude brought about by linearly equating the frequency difference between standards and deviants. Tones that are 150 Hz distal from a 600 Hz standard are 2.56 semitones more distant from their standard, compared with tones that are 150 Hz discrepant from a 1400 Hz standard. This is because semitones are measured in ratios, not simple addition. Similarly, the lowest deviant tones within each set (e.g. 450 Hz) were 0.66 semitones more distant from their standard (e.g. 600 Hz) than were the higher-pitched tone in each set (e.g. 750 Hz). Experiment 2 was sufficiently powered to detect changes in MMN magnitude brought about by both kinds of physical difference, even when those physical differences were quite small. Although it is more typical to use manipulations of several semitones (e.g. 4 to 20 semitones in May et al, 1999), MMN responses can be detected for standard-to-deviant differences of 0.27 semitones, which is near the perceptual threshold (Sams et al, 1985). The smallest of our incidental manipulations (0.66 semitones) is about twice this size, and yet we reliably detected its presence. On its own, the effect of physical deviance on MMN magnitude is unsurprising, as MMN magnitudes are well-known to be a product of deviations in frequency, duration, or intensity (see Näätänen et al., 1989; Näätänen et al., 2007; Schröger, 1998). However, this effect of physical deviance constitutes an effective ceiling for any undetected effect of reinforcement. That is, any undetected effect of reinforcement on MMN magnitude is unlikely to exceed the effect induced by a change of 0.66 semitones in the deviant tone, precisely because we reliably detected this effect.

The present null finding is somewhat surprising given that Demeter et al. (2022) recently found that high reward deviant tones elicit larger N1, MMN, and P3b compared with low-reward auditory tones. However, in their task, the critical tone stimuli were task relevant response targets, so were presumably the focus of endogenous, sustained attention. Additionally, the N1 and MMN components in their study overlapped and therefore, despite there being a functional distinction between them, they were unable to differentiate these components in their statistical analysis. Based on our results, it appears that any broad impact of associative value on electrophysiological components (including the MMN) may be limited to instances in which the response-evoking stimuli were already the subject of sustained, top-down attention, with only the P3a remaining sensitive to unexpected, significant stimuli in an unattended stream.

In addition to our ERP findings, we observed no difference in frontocentral theta coherence between conditions based on value. While we replicated earlier observations of increases in phase-locking for deviants compared with standards in both experiments (Klimesch, 1999; Rizzuto et al., 2003), we saw no modulation of this effect by associative value and Bayes factors suggested moderate evidence for a null effect of value on phase-locking in the theta range. This is notable since theta-band activity has been argued to be a particularly important spectral constituent of the auditory MMN response (Javitt et al., 2018).

#### 4.1.3. Value Impacts Behaviour and the P3a

The final reason we believe our study was sufficiently sensitive is that we observed effects of the value manipulation measured during the MMN sequence, just not on the MMN response itself. During the initial blocks of Experiment 2, participants responded slower on the visual detection task when valued deviants were presented compared with non-valued deviants. This suggests that participants knowledge of the value pairings was sufficient to interfere with their performance on the vigilance task performed during the MMN sequence. Additionally, there was indicative evidence that stimulus value may have impacted P3a magnitude. While there was no effect of value on the P3a at the whole sample level, an effect of value was observed in those participants who we might expect to show the largest impact of the value manipulation—that is, participants who demonstrated good contingency knowledge at test. Collectively, then, these results suggest that, not only were participants aware of the reinforcement contingencies, but that these contingencies influenced their behavioural and electrophysiological responses during the MMN sequence.

The observation of value modulation of the P3a to task irrelevant stimuli elicited in top learners supports earlier work investigating the P3 response to task relevant deviant stimuli. Delplanque et al. (2006) used a visual oddball paradigm and found that a posterior P3 response was enhanced for targets that were negatively valenced compared with targets that were either neutral or positively valenced, whereas a fronto-central P3 was reduced for negatively valenced targets compared with positive targets. They categorized the former response as a P3a and the latter as a P3b. However, in Delplanque et al.’s (2006) task, people were actively seeking deviant stimuli, so all response-evoking stimuli were relevant to the primary task. In our present task, all deviant stimuli were presented in an explicitly task irrelevant stream and did not require any response. In this sense, our task is more aligned with the kinds of value-driven attentional capture seen for unattended or irrelevant stimuli reported in the behavioural literature (Le Pelley et al, 2015; Anderson et al, 2016).

### 4.2. Predictive Coding Account of the MMN

Predictive coding models are arguably the dominant theory of MMN function and neurocognitive organization more generally (for review, see Spratling, 2015). Predictive coding refers to a hierarchical framework in which higher-level predictions are passed from upper layers of the cortex to lower layers, while prediction errors are passed from lower, data-driven layers to higher, more abstract layers (Friston & Kiebel, 2009; Bastos et al, 2012). A “prediction error” signal is generated within each layer whenever an expectation does not match the incoming, bottom-up sensory input. The goal of each layer is to adjust the signal weighting to minimize prediction error, and in so doing, better align the internal representation with experience. Within this framework, the MMN is characterised as a hallmark index of prediction error (Garrido et al, 2009; Wacongne, Changeux & Dehaene, 2012).

One possibility is that predictive coding models would predict the reward-pairing manipulations used in the current study to have no impact on MMN magnitudes, as the reward stimuli themselves were not present during the presentation of the tones in the MMN sequence. Since higher order beliefs about reward would not shape expectations regarding the physical properties of the upcoming tone, they ought not to have impacted a lower-level prediction error signal such as the MMN. Indeed, many observations regarding MMN modulation by higher-order expectations pertain to the physical properties of the upcoming stimulus, such as through longer-term higher-order patterns (e.g. continuously increasing Shepard tones, Tervaniemi, Maury & Naatanen, 1994), global order effects (the primacy bias; Todd et al, 2011) or the use of paired deviant tones (Todd et al., 2014).

One challenge for this interpretation is that MMN can be influenced by higher-order patterns that do not predict the physical features of the next stimulus in sequence. Mullens et al. (2014) observed that, if participants first completed a go / no-go task using standard and deviant tones, then the pattern of observed MMNs following the go / no-go task was affected by each tone’s status (i.e., whether it was a ‘go’ or ‘no-go’ stimulus). The effect was not a simple facilitation or impairment of the MMN, but instead was only evident in the degree to which order effects (the primacy bias) impacted MMN magnitude—and even then, the effect was primarily present in a single of the counterbalancing order conditions. It is also noteworthy that this primary effect modulation was itself ameliorated by the presence of a concurrent, secondary task during presentation of the auditory tone sequence (Frost et al, 2018). Ultimately, it remains possible that – like the primary bias – reward pairing might also lead to similar, second-order impacts on MMN magnitude that reflect a change in the MMNs generated for a given tone stimulus as it shifts between a high-frequency standard and rare deviant in different blocks. We were unable to test for this since tones did not alternate between standards and deviants in the present protocol, and because we used a concurrent vigilance task similar to those that remove second-order influences of abstract task knowledge on MMN.

More generally, this sensitivity of the MMN response to associative pairings with reward is interesting in the context of psychotic illness. There are several theories of psychosis that fall within the predictive framework (Fletcher & Frith, 2009; Corlett, Honey & Fletcher, 2019). While they differ in detail, each describe how sensory information may be under- or over-weighted relative to the inputs from higher-order layers, ultimately resulting in erroneous beliefs (delusions) or sensory experiences (hallucinations). Support for such accounts includes the observation that people with schizophrenia show disruptions in prediction error signalling when processing associative reward (Corlett et al., 2007; Powers et al., 2017; Waltz et al., 2007) and, separately, in processing sensory prediction errors, such as the MMN (for review, see Umbricht & Krljes, 2005). On its face, the present data would tend to challenge any simple account in which a single prediction error event is elicited through a merging of higher-order expectations about what a stimulus will look like (or how it will sound) and what it means. Instead, it appears that sensory prediction errors (i.e., the MMN) and reward prediction errors are indexed by different, but presumably temporally concurrent responses.

While there are existing EEG markers of reward processing (e.g., the Feedback Related Negativity), these are typically measured at the onset of reward or its surprising absence (for review, see Fassbender et al., 2023). In our task, reward was not specifically shown during training and instead the cueing stimulus signalled the presence or absence of reward alone. Several recent studies using different experimental tasks have all observed alteration in P300 responses to cueing stimuli when those stimuli were presented within rewarded tasks (Palidis & Gribble, 2020; Liu et al, 2020; Hughes, Mathan & Young, 2013). For this reason, we suspect that the P3a response in the present data may perform the function of a reward prediction error signal, but this conclusion requires further investigation.

## 5. Conclusion

The current study found that MMN for unattended, task irrelevant stimuli in the auditory or visual domains was insensitive to learned value associations. This conclusion held when exploratory mass univariate analyses were deployed, and when the data were separated by frequency band. This suggests that associative value does not impact upon the pre-attentive selection processes indexed by the MMN response when the response-evoking stimuli are task irrelevant. This finding contrasts with Demeter et al.’s (2022) recent observation that stimulus value affects MMN magnitude for task relevant, attended stimuli. Although stimulus value did not impact the MMN, we observed a significant effect of value on the P3a response in those participants who demonstrably knew the stimulus-mappings. Further, performance on a concurrent psychomotor vigilance task was impacted by the value of unattended stimuli. Cumulatively, these findings suggest that the P3a may index the capacity of motivationally significant stimuli to capture attention even when presented in an unattended stream (see Horvath et al., 2008). Future work will be needed to clarify the generalisability of this finding, including whether it holds in other contexts, such as for other auditory stimuli or within vision.

## Notes

### Competing Interest Statement

The authors have declared no competing interest.

